# Lymphopenic community acquired pneumonia, an unnoticed phenotype associated to mortality in non immuno-suppressed patients: a retrospective cohort study

**DOI:** 10.1101/170530

**Authors:** Jesus F Bermejo-Martin, Catia Cilloniz, Raul Mendez, Raquel Almansa, Albert Gabarrus, Adrian Ceccato, Antoni Torres, Rosario Menendez, for the NEUMONAC group

**Author notes:** Equal contribution (Catia Cilloniz, Raul Mendez, Raquel Almansa). Equal contribution (Antoni Torres, Rosario Menendez). **Researchers of the NEUMONAC group:** Pedro Pablo España. (Hospital de Galdakao, Galdakao); Luis Borderías (Hospital San Jorge, Huesca); Olga Rajas (Hospital La Princesa, Madrid); Jordi Almirall (Hospital de Mataró, Mataró); Rafael Zalacaín (Hospital de Cruces, Bilbao); Montserrat Vendrell (Hospital Josep Trueta, Girona); Salvador Bello (Hospital Miguel Servet, Zaragoza); Isabel Mir (Hospital Son Llàtzer, Palma de Mallorca); Concepción Morales (Hospital Virgen de las Nieves, Granada); Luis Molinos (Hospital Universitario Central de Asturias, Oviedo); Ricard Ferrer (Hospital Mutua Terrasa, Terrasa); Mª Luisa Briones (Hospital Clínico Universitario, Valencia); Rosa Malo (Hospital Puerta de Hierro, Majadahonda). **Corresponding author:** Antonio Torres, Department of Pneumology, Institut Clinic de Respiratori, Hospital Clinic of Barcelona-Institut d’Investigacions Biomèdiques August Pi i Sunyer (IDIBAPS), University of Barcelona (UB)-SGR 911-Ciber de Enfermedades Respiratorias (Ciberes, CB06/06/0028) Villarroel, Barcelona, Spain.

## Abstract

**Background:** The role of neutrophil and lymphocyte counts as predictors of prognosis in Community Acquired Pneumonia (CAP) has not been appropriately studied.

**Methods:** This was a retrospective study to evaluate by multivariate regression analysis, the association between neutrophil and lymphocyte counts with mortality at 30-days post discharge in two large cohorts of hospitalized patients with CAP and no prior immunosupression: a multicentric with 1550 patients recruited at 14 hospitals in Spain and a unicentric with 2840 patients recruited at the Hospital Clinic-Barcelona.

**Findings:** The unicentric cohort accounted with a higher proportion of critically ill patients: 586 (20·6%) vs 131 (8·5%) and non survivors 245 (8·6%) vs 74 (4·8%). Lymphopenia (< 1000 lymphocytes/mm^3^) was present in the 52·8% of the patients in both cohorts. A sub-group of lymphopenic patients, those with lymphocyte counts below decil 3 (677 lymphocytes/mm^3^ in the multicentric cohort and 651 lymphocytes/mm^3^ in the unicentric one), showed > 2-fold increase in the risk of mortality, independently of the CURB-65 score, critical illness and receiving an appropriated antibiotic treatment: (OR [CI95%], *p*) (2·18 [1·21- 3·92], 0·009) and (2·33 [1·61-3·33], <0·001) respectively. Neutrophil counts were not associated with mortality risk.

**Interpretation:** Lymphopenia is present in a half of the patients with CAP needing of hospitalization, in absence of antecendents of immunosupression. Lymphopenic CAP with lymphocyte counts < 664 lymphocytes/mm^3^ constitutes a particular immunological phenotype of the disease which is associated to an increased risk of mortality.

**Funding:** CibeRes, 2009 Support to Research Groups of Catalonia 911, IDIBAPS, SEPAR, SVN

## Introduction

Community-acquired pneumonia (CAP) is a serious health problem causing high morbidity and mortality worldwide ^1^. The rates of hospitalized patients due to CAP are increasing, with 22–42% of adults needing of admission to the hospital. CAP has an associated mortality of 5–14%. Around 5% of the patients hospitalized with CAP require admission to an intensive care unit (ICU). In these severe cases mortality raises to 35% ^2^.

Host individual variability is now recognized as a key factor influencing clinical expression and prognosis of CAP ^3^. Learning from the successes of utilizing precision medicine in the treatment of cancer, identifying individual phenotypes associated to poor outcome in CAP could help to understand better the pathogenesis of this disease and to improve its treatment ^4^ ^5^ ^6^. In this regard, Davenport *et al,* using a transcriptomic analysis, found a gene expression signature (SRS1) which identifies individuals with an immunosuppressed phenotype and higher 14 day mortality within a cohort of patients with sepsis secondary to CAP ^7^. In addition, new biomarkers like expression of HLADR on monocytes ^8^, soluble CD14 (presepsin) ^9^, interleukins ^10^ ^11^, procalcitonin ^12^ and mid-regional pro-ADM ^13^ could help risk assessment in the early moments of the disease. Alternatively to these new sophisticated (and expensive) “omics” and biomarker based approaches, a simple leukogram has shown to be a potential tool for classifying patients with CAP or sepsis based upon their outcome. The neutrophil/lymphocyte ratio (NLR) has been proposed as a candidate predictor of mortality for hospitalized CAP patients ^14^. In turn, we have demonstrated that septic shock patients who fail to expand circulating neutrophil counts in blood present an increased risk of mortality ^15^.

Treatment options in CAP are still reduced to antibiotics and organ support in the most severe cases ^16^. The important burden of morbidity and mortality associated to CAP has prompted investigations into a wide range of potential adjunctive immunological therapies ^16^. Some of these therapies pretend to expand neutrophil counts ^17^ ^18^, while others are addressed to increase T cell counts and/or stimulate their function ^19^ ^20^ ^21^. In our view, prior to the development of any further clinical trial with drugs expanding neutrophils or lymphocyte counts in blood, it would be necessary to elucidate whether or not they have any influence on the outcome of CAP patients.

The objective of the present study was to evaluate the impact of neutrophil and lymphocyte counts in blood on the mortality risk of patients with CAP and no antecedents of immunosupression. For this purpose, we performed a retrospective study in two large cohorts of hospitalized patients with this disease. Results of this work could help to clarify the potential role of neutrophil and lymphocyte counts on the development of precision medicine in CAP.

## Methods

### 1. Study design

Data from two different cohorts of hospitalized patients with CAP (one recruited in the context of a multicentric study and the other in a unicentric study), were employed to evaluate in a retrospective manner, the association between neutrophil and lymphocyte counts at hospital admission with the risk of mortality at 30 days following discharge. Only those patients showing complete data on lymphocyte and neutrophil counts in the first 24 hours following admission to the hospital and the variables “admission to the ward/ICU”, and “mortality at 30 days post-discharge from hospital” were considered in the analysis.

### 2. Patient Selection, Inclusion and Exclusion Criteria

#### Multi-centric cohort

The patients included in this cohort belonged to 14 Spanish hospitals (NEUMONAC group), other than Hospital Clinic (Barcelona). Patients were recruited from January 2012 to June 2015. Inclusion criteria were: presence of new pulmonary infiltrate in chest radiograph and respiratory signs and symptoms compatible with CAP (cough, expectoration, chest pain, dyspnea, fever, etc). Exclusion criteria were as follows: nursing-home patients, immunosuppresion status (human immunodeficiency virus-positive, haemathological disease, solid-organ transplantation, >14 days of treatment with >20 mg/day of prednisone or equivalent and other immunosuppresive drugs).

#### Unicentric cohort

It was constituted by consecutive patients admitted to Hospital Clinic, Barcelona, Spain, between January 2005 and December 2015 with a diagnosis of community-acquired pneumonia. Pneumonia was defined as a new pulmonary infiltrate found on the hospital admission chest radiograph and symptoms and signs of lower respiratory tract infection. Patients with prior immunosupression were excluded (e.g., patients with neutropenia after chemotherapy or bone marrow transplantation, patients with drug-induced immunosuppression as a result of solid-organ transplantation, corticosteroid (>10mg/day) or cytotoxic therapy, HIV-infected patients).

#### Other definitions

In both studies, ARDS was identified in the first 24 hours after hospital admission by applying the criteria described in the Berlin definition ^22^. Appropriateness of empiric antibiotic treatment was defined according to multidisciplinary guidelines for the management of CAP ^23^. Acute renal failure was defined using the criteria developed by the Second International Consensus Conference of the Acute Dialysis Quality Initiative (ADQI) Group ^24^. Septic shock was defined (according to the definition proposed by the American College of Chest Physicians/Society of Critical Care Medicine Consensus Conference ^25^.

### 3. Ethics

For the multicentric cohort, the Ethics Committee of the coordinating center approved the study (Code: 2011/0512). For the unicentric cohort, the Ethics Committee of the Hospital Clinic of Barcelona approved the study (Code: 2009/5451). Given to the observational and retrospective nature of the study, the informed consent was waived.

### 4. Leukocyte, lymphocyte and neutrophil quantification

This was performed on blood collected in EDTA tubes by using the automatic analyzers available in each participating hospital at the central laboratories, using standard operative procedures approved for clinical use. Lymphopenia was considered as a total lymphocyte count less than 1000 /mm^3^, following the definition proposed by Vasu S and Caligiuri MA in *Hematology*, 9th edn ^26^.

### 5. Statistical Analysis (supplementary file 1)

The association between lymphocytes / neutrophils and the risk of mortality at 30 days following hospital discharge was assessed by multivariate logistic regression analysis. Potential confounding variables were selected by assessing the association between those variables showed in supplementary file 1 with mortality. Variables yielding a *p* < 0·1 in the univariate regression analysis were further included in the multivariate one as adjusting variables. Final selection of the variables was performed by using the backward stepwise selection method (Likehood Ratio) [pin<0·05, pout<0·10]). Lymphocyte and neutrophil concentrations in blood were transformed to Naeperian log values in order to reach a normal distribution. Those variables showing a Spearman correlation coefficient > 0·3 with another inclusive variable were excluded from the multivariate analysis. Final “n” for each analysis (excluding the missing values) is showed in tables 1 to 5. Data were analyzed by using the IBM SPSS 20·0 software (SPSS, Chicago, Ill).

### 6. Role of the funding source

The institutions supporting this work did not have any role in the in study design, in the collection, analysis, and interpretation of data, in the writing of the report or in the decision to submit the paper for publication.

## Results

### Clinical characteristics of the patients (table 1)

The multicentric cohort accounted with1550 patients, for 2840 of the unicentric cohort. The proportion of patients older than 65 *yr* was higher in the multicentric study. In turn, the proportion of those who had received vaccination against *S. pneumoniae* or *influenza virus* was higher in the unicentric study. Prevalence of chronic pulmonary disease and neurological disease was also higher in those patients recruited in the unicentric cohort. This cohort was in turn more severe, as evidenced by the increased proportion of patients showing a CURB-65 score ≥ 3, of patients deserving admission to the ICU and finally of patients showing renal failure, pleural effusion, septic shock and mechanical ventilation as complications during hospitalization. In agreement with this scenario, the frequency of non survivors in the unicentric cohort was almost two-fold than that in the multicentric one, although the majority of the patients in this cohort had received an appropriated antibiotic treatment. Patients in the multicentric cohort showed more frequently an antecedent of cardiac disease. Patients in this cohort showed a higher proportion of pneumococcal CAP cases and lower proportion of viral CAP within those patients with a positive microbiological identification. Length of hospitalization was 6 (4-9) days in the multicentric study and 8 days (6-12) in the unicentric one (median, interquartilic range). The percentage of patients showing lymphopenia in each cohort was the same, 52·8% (818 out of 1550 patients in the multicentric study and 1499 patients out of 2840 patients in the unicentric study). Leukocyte counts were similar in both cohorts: Leucocytes (cells/mm3): 13000 (9200-17790)/ 12300 (8400-17300); Neutrophils (cells/mm3): 10823 (7274-15187) / 10126 (6636-14700); Lymphocytes (cells/mm3): 957 (616-1445) / 950 (578-1440) in the multicentric study and unicentric study respectively [median (interquartile range)].

**Table 1:**
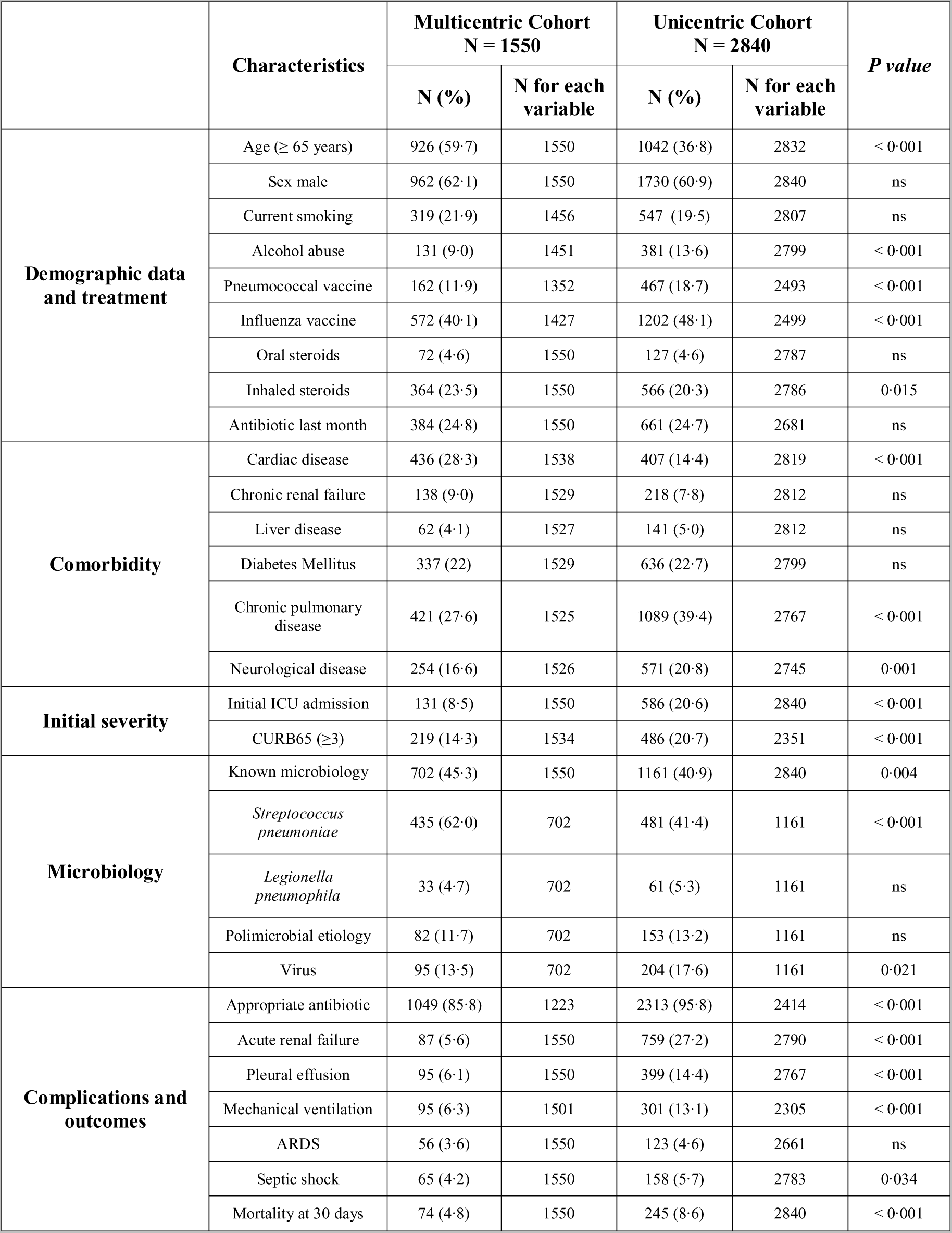
Clinical characteristics of the studied cohorts. Data presented as n (%) or median (interquartile range). Microbiological data were calculated using the number of patients with positive microbiological identification. ICU: Intensive care unit, ARDS: Acute Respiratory Distress Syndrome.

### Analysis of mortality risk at 30 days following hospital discharge

#### Multi-centric cohort

In the univariate analysis, the lymphocyte count was inversely associated to the risk of mortality (*p* < 0·05, Table 2). In contrast, in this analysis neutrophils showed no significant association with mortality: (0·77 [0·53 – 1·11], *p* = 0·16) (OR [CI95%], *p*). Because of that, the multivariate analysis was focused on the lymphocyte count. A complete description of the uni and multivariate analysis process is detailed in the Supplementary figure 1. The multivariate analysis confirmed that the lymphocyte count was a protective factor against mortality, independently of the variables [presence of chronic renal failure], [CURB-65 score], [initial admission to the ICU], and also of [appropriate antibiotic treatment] (Table 2). When the multivariate analysis was repeated distributing the patients in categories based on lymphocyte decils, the highest decil showing a significant association with mortality risk was 677 lymphocytes/mm^3^ (decil 3) (Table 3). This way, showing < 677 lymphocytes/mm^3^ increased by 2·18 the probability of dying in the 30 days following discharge from hospital (Table 3). 8·4% of the patients in the group with < 677 lymphocytes/mm^3^ have died by day 30 following hospital admission, for 3·2% in the group with ≥ 677 lymphocytes/mm^3^ (Supp table 1). Patients with < 677 lymphocytes/mm^3^ presented more frequently with critical illness (Supp table 1). This group showed an increased proportion of patients with ≥ 3 points in the CURB-65 score, along with a higher frequency of complications (pleural effusion, mechanical ventilation, ARDS, septic shock) (Supp table 1). Patients with < 677 lymphocytes/mm^3^ had more frequently a positive microbiological identification.

**Table 2:**
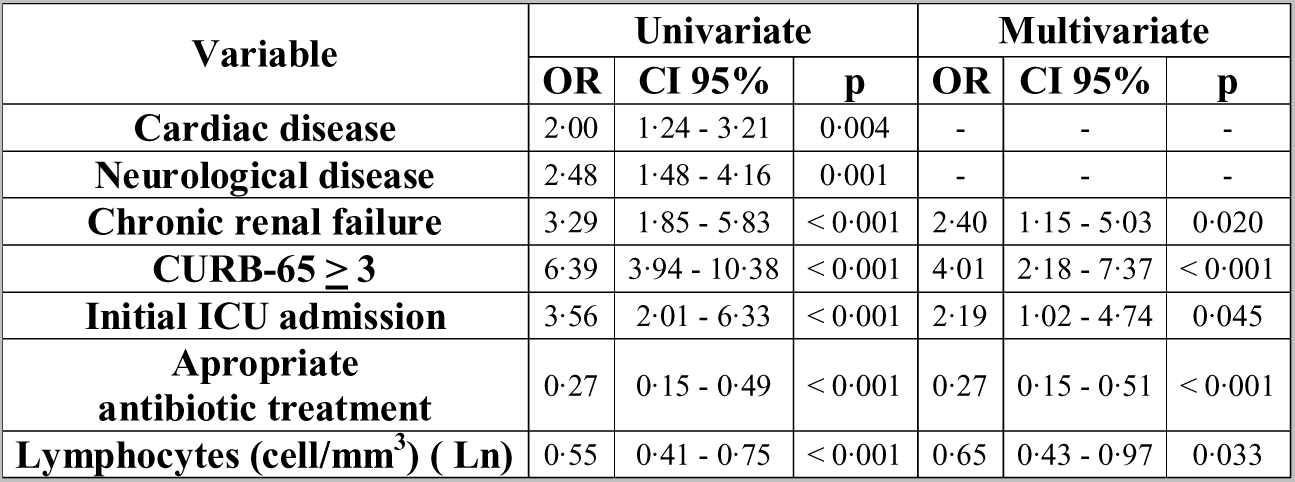
Univariate and Multivariate regression analysis for mortality risk in the multicentric cohort. OR: Odds Ratio; CI: confidence interval. Total “n” of patients included in the the multivariate analysis was 1209.

| Variable | Univariate |  |  | Multivariate |  |  |
| --- | --- | --- | --- | --- | --- | --- |
|  | OR | CI 95% | p | OR | CI 95% | p |
| **Cardiac disease** | 2.00 | 1.24 - 3.21 | 0.004 | - | - | - |
| **Neurological disease** | 2.48 | 1.48 - 4.16 | 0.001 | - | - | - |
| **Chronic renal failure** | 3.29 | 1.85 - 5.83 | < 0.001 | 2.40 | 1.15 - 5.03 | 0.020 |
| **CURB-65 ≥ 3** | 6.39 | 3.94 - 10.38 | < 0.001 | 4.01 | 2.18 - 7.37 | < 0.001 |
| **Initial ICU admission** | 3.56 | 2.01 - 6.33 | < 0.001 | 2.19 | 1.02 - 4.74 | 0.045 |
| **Aproprate antibiotic treatment** | 0.27 | 0.15 - 0.49 | < 0.001 | 0.27 | 0.15 - 0.51 | < 0.001 |
| **Lymphocytes (cell/mm<sup>3</sup>) ( Ln)** | 0.55 | 0.41 - 0.75 | < 0.001 | 0.65 | 0.43 - 0.97 | 0.033 |

**Table 3:** Multivariate regression analysis for mortality risk by decils of lymphocyte counts in the multicentric cohort. OR: Odds Ratio; CI: confidence interval. Total “n” of patients included in the the multivariate analysis was 1209.

|  | Analysis for decil 1<br>( < 414 lymphocyte/mm<sup>3</sup> ) |  |  | Analysis for decil 2<br>( < 551 lymphocyte/mm<sup>3</sup> ) |  |  | Analysis for decil 3<br>( < 677 lymphocyte/mm<sup>3</sup> ) |  |  |
| --- | --- | --- | --- | --- | --- | --- | --- | --- | --- |
| Variable | OR | CI 95% | *p* | OR | CI 95% | *p* | OR | CI 95% | *p* |
| Chronic renal failure | 2.23 | 1.04 - 4.74 | 0.038 | 2.29 | 1.09 - 4.84 | 0.029 | 2.38 | 1.13 – 5.00 | 0.022 |
| CURB-65 ≥ 3 | 3.64 | 1.95 - 6.77 | < 0.001 | 3.90 | 2.11 - 7.21 | < 0.001 | 3.89 | 2.11 - 7.19 | < 0.001 |
| Initial ICU admission | 2.21 | 1.02 - 4.78 | 0.044 | 2.11 | 0.98 - 4.54 | 0.057 | 2.16 | 1.00 - 4.64 | 0.049 |
| Aproprate antibiotic treatment | 0.27 | 0.15 - 0.51 | < 0.001 | 0.27 | 0.15 - 0.50 | < 0.001 | 0.27 | 0.15 - 0.50 | < 0.001 |
| [Lymphocytes] | 4.09 | 2.13 – 7.85 | < 0.001 | 2.83 | 1.56- 5.13 | 0.001 | 2.18 | 1.21- 3.92 | 0.009 |

#### Uni-centric cohort

As occurred in the multicentric cohort, in the univariate analysis the lymphocyte count was a protective factor against mortality (*p* < 0·05, Table 4), with neutrophils failing to show any significant results in this analysis (0·95 [0·88 – 1·01], *p* = 0·11) (OR [CI95%], *p*). The multivariate analysis confirmed that the lymphocyte count was a protective factor against mortality, independently of the variables [presence of neurological disease], [CURB-65 score], [initial admission to the ICU], [development of ARDS as complication during hospitalization], and [appropriate antibiotic treatment] (Table 4). Variables included in the uni and multivariate analysis are detailed in the Supplementary figure 1. Similarly to that found in the multicentric cohort, showing lymphocyte counts below decil 3 (651 lymphocytes / mm^3^) translated into a 2·33 fold increase in the risk of mortality (Table 5). 12·9 % of the patients in the group with < 651 lymphocytes/mm^3^ have died by day 30 following hospital admission, for 6·8 % in the group with ≥ 651 lymphocytes/mm^3^ (Supp table 2). Patients with < 651 lymphocytes/mm^3^ presented more frequently with critical illness (Supp table 2). This group showed a higher frequency of complications (acute renal failure, mechanical ventilation and septic shock) (Supp table 2). Patients with < 651 lymphocytes/mm^3^ had more frequently a positive microbiological identification (Supp table 2). In this cohort, the highest decil showing a significant association with mortality risk was decil 5 (950 lymphocytes/mm^3^) (Table 5).

**Table 4:**
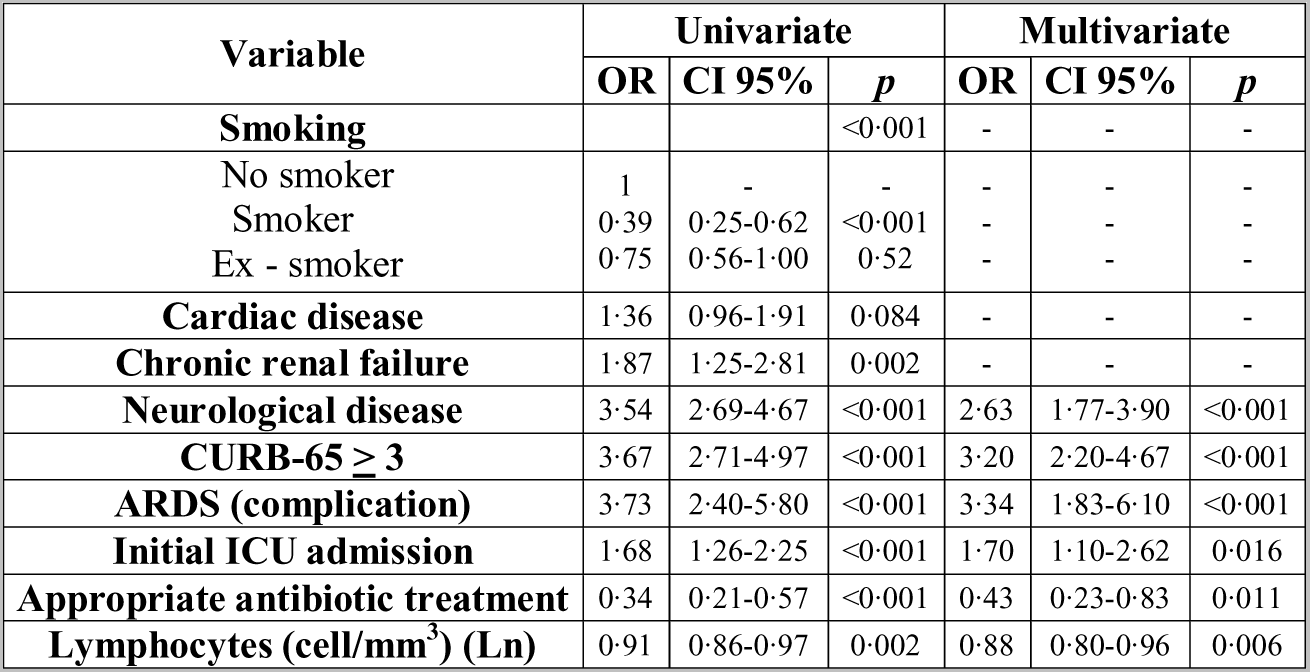
Univariate and Multivariate regression analysis for mortality risk in the unicentric cohort. OR: Odds Ratio; CI: confidence interval. Total “n” of patients included in the multivariate analysis was 1851.

| Variable | Univariate |  |  | Multivariate |  |  |
| --- | --- | --- | --- | --- | --- | --- |
|  | OR | CI 95% | *p* | OR | CI 95% | *p* |
| **Smoking** |  |  | <0.001 | - | - | - |
| No smoker | 1 | - | - | - | - | - |
| Smoker | 0.39 | 0.25-0.62 | <0.001 | - | - | - |
| Ex - smoker | 0.75 | 0.56-1.00 | 0.52 | - | - | - |
| **Cardiac disease** | 1.36 | 0.96-1.91 | 0.084 | - | - | - |
| **Chronic renal failure** | 1.87 | 1.25-2.81 | 0.002 | - | - | - |
| **Neurological disease** | 3.54 | 2.69-4.67 | <0.001 | 2.63 | 1.77-3.90 | <0.001 |
| **CURB-65 $\geq 3$** | 3.67 | 2.71-4.97 | <0.001 | 3.20 | 2.20-4.67 | <0.001 |
| **ARDS (complication)** | 3.73 | 2.40-5.80 | <0.001 | 3.34 | 1.83-6.10 | <0.001 |
| **Initial ICU admission** | 1.68 | 1.26-2.25 | <0.001 | 1.70 | 1.10-2.62 | 0.016 |
| **Appropriate antibiotic treatment** | 0.34 | 0.21-0.57 | <0.001 | 0.43 | 0.23-0.83 | 0.011 |
| **Lymphocytes (cell/mm<sup>3</sup>) (Ln)** | 0.91 | 0.86-0.97 | 0.002 | 0.88 | 0.80-0.96 | 0.006 |

**Table 5:** Multivariate regression analysis for mortality risk by decils of lymphocyte counts in the unicentric cohort. OR: Odds Ratio; CI: confidence interval. Total “n” of patients included in the multivariate analysis was 1851.

|  | Analysis for decil 1<br>( < 295 lymphocyte/mm<sup>3</sup>) |  |  | Analysis for decil 2<br>( < 500 lymphocyte/mm<sup>3</sup>) |  |  | Analysis for decil 3<br>( < 651 lymphocyte/mm<sup>3</sup>) |  |  | Analysis for decil 4<br>( < 780 lymphocyte/mm<sup>3</sup>) |  |  | Analysis for decil 5<br>( < 950 lymphocyte/mm<sup>3</sup>) |  |  |
| --- | --- | --- | --- | --- | --- | --- | --- | --- | --- | --- | --- | --- | --- | --- | --- |
| Variable | OR | CI 95% | *p* | OR | CI 95% | *p* | OR | CI 95% | *p* | OR | CI 95% | *p* | OR | CI 95% | *p* |
| Neurological disease | 2.64 | 1.78-3.92 | <0.001 | 2.62 | 1.77-3.87 | <0.001 | 2.60 | 1.76-3.84 | <0.001 | 2.62 | 1.77-3.86 | <0.001 | 2.62 | 1.78-3.87 | <0.001 |
| CURB-65 ≥ 3 | 3.11 | 2.13-4.53 | <0.001 | 3.06 | 2.10-4.45 | <0.001 | 3.09 | 2.13-4.50 | <0.001 | 3.07 | 2.12-4.45 | <0.001 | 3.09 | 2.13-4.48 | <0.001 |
| ARDS (complication) | 3.29 | 1.79-6.04 | <0.001 | 3.09 | 1.68-5.69 | <0.001 | 3.39 | 1.85-6.24 | <0.001 | 3.24 | 1.77-5.92 | <0.001 | 3.21 | 1.76-5.85 | <0.001 |
| Initial ICU admission | 1.67 | 1.08-2.58 | 0.021 | 1.46 | 0.94-2.27 | 0.089 | 1.48 | 0.96-2.28 | 0.080 | 1.53 | 0.99-2.36 | 0.054 | 1.55 | 1.01-2.39 | 0.047 |
| Appropriate antibiotic treatment | 0.43 | 0.22-0.81 | 0.010 | 0.40 | 0.21-0.76 | 0.005 | 0.44 | 0.23-0.83 | 0.012 | 0.44 | 0.23-0.84 | 0.013 | 0.43 | 0.23-0.81 | 0.010 |
| [Lymphocytes] | 2.33 | 1.37-3.85 | 0.001 | 2.44 | 1.64-3.57 | <0.001 | 2.33 | 1.61-3.33 | <0.001 | 1.75 | 1.20-2.50 | 0.003 | 1.75 | 1.20-2.56 | 0.003 |

## Discussion

Our study revealed a first surprising finding: 52·8% of the patients with CAP show lymphopenia at hospital admission (< 1000 lymphocytes/mm^3^) ^26^, in absence of antecedents of immunosupression. Revisiting the data of another large cohort from Marrie TJ and Wu L, including 3043 hospitalized patients with CAP and no antecedents of immunsupression^27^, a similar prevalence of lymphopenia was observed at hospital admission (48·4%). In our view, the high prevalence of lymphopenia in CAP has gone largely unnoticed until the present moment. Lymphopenia could be a cause or a consequence of CAP. Impaired production or increased apoptosis of lymphocytes caused by the presence of chronic diseases or critical illness ^28^, enhanced adhesion to the vascular endothelium or massive migration of these cells to the lungs could explain the presence of lymphopenia in some patients with this disease.

Interestingly, we identified a sub-group of lymphopenic patients, those with lymphocyte counts below decil 3 (651 lymphocytes/mm^3^ in the unicentric cohort and 677 lymphocytes/mm^3^ in the multicentric one), which showed a 2-fold increase in the risk of mortality. This subgroup accounted with an increased proportion of critically ill patients and of those who developed complications (supplementary table 1 and 2). These patients had also a higher frequency of positive microbiological identification (supplementary table 1 and 2). If this is consequence of a higher microbial load secondary to a poor control of infection in these patients deserves to be investigated in future works. In consequence, our findings suggest that CAP patients with lymphocyte counts below decil 3 constitute a particular immunological phenotype of the disease which is associated with the presence of a more severe disease and worst outcome.

In the work from Marrie TJ and Wu L, a lymphocyte count < 1000 cell/mm^3^ count was associated with early but not with late mortality. Nonetheless, there are a number of important differences between this work and ours. The most important is that Marrie TJ and Wu L excluded patients with leukocyte counts < 1000 cells/mm^3^ and those with critical illness. In addition, they did not evaluate lymphocyte counts as a continuous variable to predict mortality, or other cut-offs below 1000 lymphocytes/mm3. Finally, they evaluated hospital mortality, not mortality at 30 days post-discharge as we did.

Early identification of patients at risk of death is a crucial step in the clinical management of the patients suffering from CAP ^6^. For clinical operationalization, the mean value between both thresholds corresponding to decil 3 (664 lymphocytes/mm^3^) could help to early recognize CAP patients at risk of poor outcome. Prompt identification of these patients could contribute to improve their prognosis, since they might benefit from more aggressive treatment strategies. In this regard, evaluation of lymphocyte counts is an inexpensive, widely available test in clinical settings worldwide.

Our results support also that lymphocyte counts should be taken into consideration as a potential confusion factor in the design and analysis of future clinical trials evaluating drugs or interventions for the treatment of CAP, since they influences prognosis. In addition, our findings suggest that patients with low lymphocyte counts could need of new therapeutic approaches, since the multivariate analysis demonstrated that the presence of low lymphocyte counts conferred an increased risk of mortality independently of receiving an appropriated antibiotic treatment and intensive care. In this regard, adjunctive therapy with drugs inducing expansion of lymphocyte counts and/or modulating function of these cells ^19^ ^20^ ^21^ could represent an option for these patients to be explored in future clinical trials, as actually occurs in sepsis: NCT00711620 (Thymosin α1), NCT02640807, NCT02797431 (IL-7) or NCT02576457 (PD-L1 inhibitor) (ClinicalTrials.gov identifier).

The cohorts evaluated in this study differed in severity. This probably explained that the the upper threshold associated to mortality in the unicentric study, although being also in the range of lymphopenia, raised to 950 lymphocytes/mm^3^. Nonetheless, the threshold corresponding to decil 3 in both cohorts was very similar, evidencing that it was a preserved predictor of mortality independently of CAP severity at hospital admission. Far from representing a disadvantage, using heterogeneous cohorts could represent an opportunity. As supports the recent success of multicohort analysis and data sharing to discover new biomarkers in organ transplant, cancer, autoimmune and infectious diseases, studying heterogeneous cohorts makes that biomarkers that are discovered in these kind of cohorts are more likely to be generalizable across a broad spectrum of patients ^29^. In this sense, our work is the first in utilizing two large cohorts and the same robust multivariate analysis to support the impact of lymphopenia on the outcome of patients with CAP.

Finally, our results showed that neutrophil counts were not associated with mortality risk in patients with CAP, raising doubts on the potential role of drugs aimed to expand neutrophils in this disease. In this sense, a systematic review from Cheng AC *et al* in 2007 concluded that the use of G-CSF as an adjunct to antibiotics was not associated with improved 28-day mortality ^30^. In addition, at the light of the absence of any significant association between neutrophil counts with mortality, the potential value of the neutrophil to lymphocyte ratio as biomarker of mortality in CAP should be carefully re-evaluated.

A limitation of our work was its retrospective nature and the absence of data on lymphocyte subsets. The specific impact of CD4, CD8 T cells, B and NK lymphocytes on mortality should be studied in future prospective studies.

## Conclusions

1. Lymphopenia is a very frequent finding in hospitalized patients with CAP and no immunosupression, being present in a half of the cases at admission. 2. Lymphopenic CAP with lymphocyte counts less than 664 lymphocytes/mm^3^ constitutes a particular immunological phenotype of the disease which is associated with a two fold increase in the risk of mortality. 3. Assessing lymphocyte counts at hospital admission could contribute to personalize clinical management and treatment in CAP.

## Contributors

JFBM, AT, Rosario M conceived the study and contributed to drafting of the paper. AG, participated in the statistical analysis. CC contributed to the study concept and design and contributed to drafting of the paper. Raúl M participated in the statistical analysis and contributed to drafting of the paper. RA contributed to the study concept and design, participated in the statistical analysis and contributed to drafting of the paper. AC: contributed to the study concept and design. The NEUMONAC group participated in the acquisition of data. All authors participated in the analysis or interpretation of data.

## Declaration of interest

The authors declare no conflicts of interest regarding this submission

## Acknowledgements

We thank the nurse teams of the participant hospitals for their help with sample collection for leukogram analysis through the years. The study was funded by Ciber de Enfermedades Respiratorias (CibeRes CB06/06/0028), 2009 Support to Research Groups of Catalonia 911, IDIBAPS, “Convocatoria extraordinaria PII’s-SEPAR 2011 ref: 1046/2011”, “Beca SEPAR 2012 ref:145/2012, and “Beca Sociedad valenciana de Neumologia (SVN) 2013”. Catia Cilloriz is recipient of ERS Short Term Fellowship and Postdoctoral Grant “Strategic plan for research and innovation in health-PERIS 2016-2020”. Raquel Almansa and Jesús F Bermejo are supported by Consejería de Sanidad de Castilla y León-IECSCYL.

## Research in context

### Evidence before this study

References for this article were identified through searches of PubMed for articles published in English before July 1^st^ 2017 by use of the terms “community acquired pneumonia” and “mortality”, concomitant with the terms “biomarkers” OR “lymphocyte” OR “neutrophil” OR “adjuvant” OR “immunomodulatory therapy” OR “lymphopenia”. The most representative articles corresponding to these items were cited and discussed. We only considered peer-reviewed, English language reports, with the exception of the Multidisciplinary guidelines for the management of community-acquired pneumonia (CAP) of the Spanish Society of Respiratory Medicine and Thoracic Surgery, which are adapted for the management of CAP in Spain, and were published in the Spanish journal “Medicina Clinica”. Two articles reviewed the epidemiology of CAP; four were focused on precision medicine and new aspects in the management of this disease, with emphasis in the necessity of identifying specific phenotypes of this condition; nine articles updated the information on biomarkers in CAP, with two of this articles reviewing the use of leukogram to predict mortality; seven articles versed on immunomodulatory adjuvant treatments in CAP and sepsis; five articles corresponded to clinical definitions (ARDS, sepsis/Septic shock, appropriateness of empiric antibiotic treatment in CAP, acute renal failure and lymphopenia); one article corresponded to the only study similar to ours found in the literature, which is discussed in our article; finally, two articles were cited to discus on the impact of critical illness and chronic diseases on the immune system and the use of heterogeneous patients `cohorts for biomarker discovery.

### Added value of this study

Using two large cohorts of patients with no immunosupression, one multicentric and another one unicentric, our study reveals in first place an unnoticed finding in CAP: the existence of lymphopenia (<1000 lymphocytes / mm^3^ in blood) in a a half of the patients needing of hospitalization due to this disease. Our work identified a subgroup of patients with lymphopenia (those with < 677 lymphocytes / mm^3^ in the multicentric study and < 651 lymphocytes / mm^3^ in the unicentric one) which presented a two-fold increase in the mortality risk at 30 days post-discharge from hospital, independently of the severity of the disease and also of receiving an appropriated antibiotic treatment.

### Implications of all the available evidence

This group of patients with lymphopenic CAP constitutes a particular immunological phenotype of the disease which is at risk of poor outcome, and could be early identified using a simple leukogram, a test easily available in hospital laboratories worldwide. This phenotype should be considered as a potential confusion factor influencing outcome in the clinical trials testing new drugs for CAP. In turn, these patients could potentially benefit of specific interventions with prompt/more aggressive treatment or also of adjuvant therapies with immunomodulators stimulating lymphocyte expansion or function.

